# Epitomic analysis of the specificity of conformation-dependent, anti-Aß amyloid monoclonal antibodies

**DOI:** 10.1101/2020.08.05.238105

**Authors:** Jorge Mauricio Reyes-Ruiz, Rie Nakajima, Ibtisam Baghallab, Luki Goldschmidt, Justyna Sosna, Phuong Nguyen Mai Ho, Taha Kumosani, Philip L. Felgner, Charles Glabe

## Abstract

Antibodies against Aß amyloid are indispensable research tools and potential therapeutics for Alzheimer’s Disease, but display several unusual properties, such as specificity for aggregated forms of the peptide, ability to distinguish polymorphic aggregate structures and ability to recognize generic aggregation-related epitopes formed by unrelated amyloid sequences. Understanding the mechanisms underlying these unusual properties of anti-amyloid antibodies and the structures of their corresponding epitopes is crucial for the understanding why antibodies display different therapeutic activities and for the development of more effective therapeutic agents. Here we employed a novel “epitomic” approach to map the fine structure of the epitopes of 28 monoclonal antibodies against amyloid-beta using immunoselection of random sequences from a phage display library, deep sequencing and pattern analysis to define the critical sequence elements recognized by the antibodies. Although most of the antibodies map to major linear epitopes in the amino terminal 1-14 residues of Aß, the antibodies display differences in the target sequence residues that are critical for binding and in their individual preferences for non-target residues, indicating that the antibodies bind to alternative conformations of the sequence by different mechanisms. Epitomic analysis also identifies more discontinuous, non-overlapping sequence Aß segments than peptide array approaches that may constitute the conformational epitopes that underlie the aggregation specificity of antibodies. Aggregation specific antibodies recognize sequences that display a significantly higher predicted propensity for forming amyloid than antibodies that recognize monomer, indicating that the ability of random sequences to aggregate into amyloid is a critical element of their binding mechanism.

## Introduction

Amyloids are intermolecularly hydrogen bonded ß-sheet aggregates that have a regularly repeating lattice structure (1,2). Unlike natively folded proteins that adopt a single or limited range of structures or states, amyloids can adopt a large number of different ß-sheet aggregate structures that vary in the parallel vs antiparallel strand arrangement, the segments of the sequence that form intermolecularly hydrogen bonded ß-sheets and the locations where the sheets fold and how the sheets stack together. Many proteins are able to form amyloids upon unfolding or misfolding and are frequently associated with neurodegenerative diseases such as Alzheimer’s disease (AD), Parkinson’s disease, amyotrophic lateral sclerosis and prion diseases (3). Alzheimer’s disease contains two canonical amyloids: Aß amyloid, derived from APP and tau amyloid and in approximately 30% of AD, cortical Lewy bodies are also present containing α-synuclein amyloid. Monoclonal antibodies against Aß are a leading class of therapeutic for AD. Many of these antibodies have been evaluated in clinical trials, but so far none have demonstrated consistent therapeutic activity in slowing the progression of AD and none have been approved by the FDA (reviewed in (4,5)).

Conformation-dependent monoclonal antibodies against Aß are also an invaluable tool for research in the role of amyloids in Alzheimer’s disease because they recognize epitopes that are differentially displayed on distinct structural polymorphs or folded states of the peptide, providing insight into the role of polymorphisms in the pathogenic spectrum of the disease (6). Many of the antibodies raised against Aß amyloid specifically bind to aggregated oligomeric or fibrillar forms of the peptide and do not bind to monomer or the amyloid precursor protein (7–10). Moreover, many of these aggregation state-specific antibodies recognize aggregates formed from unrelated sequences, indicating that amyloids display generic epitopes as a consequence of their aggregated structure. Understanding the mechanisms underlying the unusual specificities of anti-amyloid antibodies and the structures of their corresponding epitopes is crucial for the understanding of the immune response to amyloid and for the development of effective therapeutic agents.

We have previously reported polyclonal rabbit sera that distinguish two classes of amyloid aggregates. A11 sera was raised against oligomeric Aß mimics made of Aß42 coupled via the carboxyl terminus to colloidal gold that recognizes prefibrillar oligomers, but not fibrils or monomer (7). Subsequently we reported OC serum that was raised against Aß42 fibrils that is specific for fibrils and fibrillar oligomers but not prefibrillar oligomers or monomer (11). A11 and OC serum recognize prefibrillar oligomers and fibrils respectively from several other amyloid forming proteins, including α-synuclein and islet amyloid polypeptide (IAPP), indicating that some of the antibodies in the sera recognize generic amyloid aggregates that do not depend on the precise peptide sequence. We also raised rabbit antiserum against Aß42 annular protofibrils, which are pore-like structures, but this serum labels the same type of prefibrillar oligomers as A11, indicating that these sera have related specificities (12). We cloned 6 monoclonals from A11 serum-producing rabbits (9), 2 monoclonals from annular protofibril vaccinated rabbits (Glabe, unpublished) and 23 monoclonals from OC serum-producing rabbits (10). For the OC monoclonals (mOC), the epitope was mapped using the PepSpots method of overlapping 10mers that vary by a single amino acid from position −5 to 45. Nineteen of the 23 mOC monoclonals gave a pattern of reactivity that mapped to either a linear or discontinuous epitope in the amino terminal 2/3 of the molecule (10) while 16 map to the amino terminal residues from 1-11 (10). Although these antibodies bind to the same regions of Aß, they bind selectively to alternative fibrils structures of Aß, indicating that this region is conformationally polymorphic (13).

Here we report that the antibodies have very different epitopes and binding modes in terms of the 1) target residues that participate in binding. 2) target residues that are not important for binding 3) non-target residues that are preferred over target residues for binding. 4) presence and location of discontinuous conformational epitopes. The significance is that this information can be used for modeling the structures of the antibody-epitope interaction and the specific peptidescan be used in an array as a fingerprint to uniquely identify antibodies that bind to these epitopes and mimotopes in complex mixtures, like human serum to identify antibodies that may be protective or predictive of disease.

## Results

### Library randomness, bias, and non-specific binding of phage

Library randomness and amino acid bias are important for distinguishing specific and non-specific binding. In order to assess the randomness and amino acid bias in the library as purchased from the manufacturer, DNA from an aliquot of the library was extracted, the random sequences amplified by PCR and deep sequenced, yielding 19,434 sequences of which 97.4% were unique sequences (Supplemental Information file 1, LibrarySeqOnly, column A, Original unamplified library). We also amplified the library 2 successive rounds and examined the effect on the number of unique sequences observed. (Supplemental Information file 1, LibrarySeqOnly, columns B, C, Amplified round 1 and round 2). In the first round of amplification, 93.4% of the sequences were unique while in the second round of amplification 93.9% were unique. To further compare the randomness of the library, we compared the unique sequences between the original library and the first amplification and found that 2.1% of the unique sequences were observed in both libraries. This is the same as the number of unique sequences that were observed twice in the same original unamplified library (2.1%). These results indicate that while some sequences may have amplified preferentially, the vast majority of sequences in the library remain random after amplification.

The library has an inherent bias in terms of the frequency of nucleotides observed as shown in Supplemental Information Table I. Thymidine is over-represented, while adenine is the lowest abundance. We also calculated the frequency of amino acid residues observed in the random library as shown in Supplemental Information Table II. Not surprisingly, leucine and serine, amino acids that have 6 codons are abundantly encountered, although arginine which also has 6 codons is encountered with approximately half the frequency. Threonine with 4 codons is also abundant, while proline is surprisingly abundant with only 2 codons. Cysteine is the lowest abundance amino acid observed. These results are very similar to the manufacturer’s observed amino acid frequency published in their data sheet for the library. Stop codons are observed with a low frequency of 0.45%, presumably because stop codons halt the expression of the pIII protein that the random sequence is fused to that is required for infectivity. While the host strain contains an amber suppressor tRNA to decode the UAG stop codon as glutamine, it is possible that other stop codons may arise from the co-transfection of a functional and non-functional pIII encoding genome into the same bacterium and packaged into phage. It is not clear why any stop codons are observed if pIII is absolutely required for infectious activity, but it is possible that this may arise from the initial transfection of the phagemid where a sequence containing a stop codon is co-transfected into the same bacterium and packaged into phage.

Non-specific binding may have a significant deleterious effect on the analysis of the patterns observed that bind to the paratope of the antigen combining site of the antibody because they do not fit any specific pattern and therefore lower the minimum percentage of the total sequences needed to match an epitope pattern (see C% below). We found that there are two types of non-specific binding sequences. The first type are random sequences that bind specifically to the protein A magnetic beads and the constant regions of the antibodies. The patterns of key amino acid residues for antibody binding that arise from these sequences are found in all replicates of the control beads without any antibody and also found in all replicates containing antibody of the same isotype regardless of antibody’s specificity. These sequences were identified and removed from the sequence files in the BASH processing script used to extract the peptide sequences from the Illumina data files. The second type of non-specific binding sequences arise from the binding of phage to the beads or antibody through an interaction other than with the random sequence fused to pIII. These sequences display the same frequency distribution as the unselected random library in terms of the presence of a high percentage of unique sequences observed only once in the sequencing run.

The purpose of repeating the immunoselection step is to increase the proportion of specific sequences, so we investigated the effect of immunoselection followed by amplification of the phage at each of the 3 steps. Using antibody mOC1 as an example, in the first immunoselection step prior to amplification 73.2% of the sequences occur a single time. At the second immunoselection prior to amplification, 98.7% are single. In the third immunoselection, 74.0% of the sequences are single reads, indicating that a substantial amount of non-specific random sequences remains even after repeated immunoselection.

### Epitomic analysis of monoclonal antibody specificity

We performed 3 immunoselection or panning steps for each antibody amplifying the eluted phage after each step and sequenced each round of panning before and after amplification as shown in Figure 1. We analyzed all 6 of these sequencing groups and found that the specific binding patterns were largely the same for each antibody, although the number of specific sequences increases with successive pannings (data not shown). We chose to present the data for the unamplified samples after 3 rounds of pannings because they have been subjected to 3 rounds of immunoselection and only 2 rounds of amplification (Supplemental Information file 2 Immunoselected sequences). A total of 6,282,154 sequence reads were obtained from the 28 monoclonals including the unamplified and amplified steps of the three pannings. This includes 22 of the mOC series monoclonals and 6 of the mA11 monoclonals. Two of the mA11 monoclonals were not analyzed because the hybridomas producing them were lost (mA11-201 and mA11-121) and the two monoclonals raised against Aß42 annular protofibrils (mA11-09 and mA11-89) were included with the mA11 monoclonals because their properties overlap in terms of binding to ß-barrel structures like annular protofibrils and hemolysins (14,15). Once the sequences were processed by removing the non-specific sequences and duplicates, we found a total of 1,713,634 unique peptide sequences. For each unique sequence found, we also counted the number of times it was observed. For the third unamplified step used in the analysis, the range was from 223 unique sequences for mOC51 to 53,621 for mOC98. The average number of unique sequences per monoclonal in the third unamplified panning was 10,959. The unique sequences were sorted by the descending number of times their sequence was observed as a surrogate for relative binding activity.

**Figure 1.**
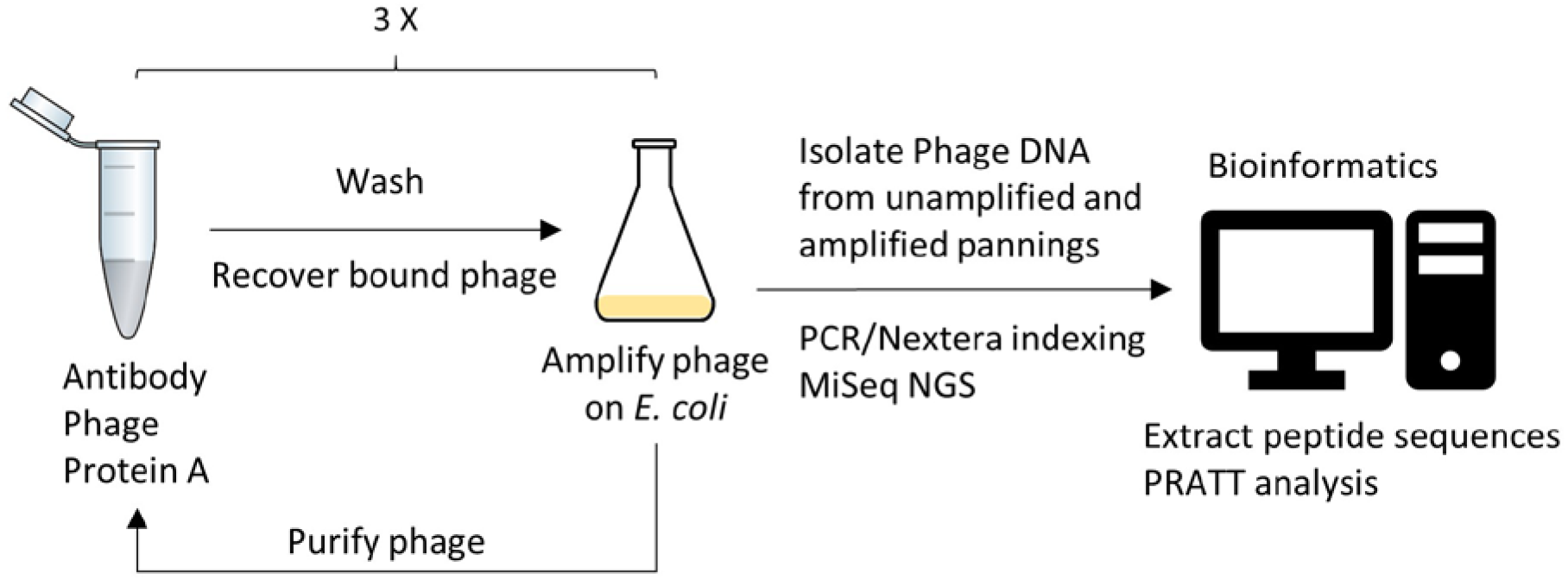
Schematic diagram for epitomic analysis of antibody specificity. Antibody is mixed with phage and Protein A beads. After washing, the phage are eluted from the beads and then the phage are amplified in *E. coli*, purified and used for antibody panning two more times. Samples of phage are taken after elution prior to amplification and after amplification. The isolated phage DNA is amplified and bar coded by PCR prior to MiSeq next generation sequencing. The peptide sequences are extracted and the unique sequences are counted and rendered in FASTA format for PRATT pattern analysis.

The sequences were analyzed using the PRATT 2.1 amino acid sequence pattern recognition algorithm that is a method for the identification of patterns in a set of unaligned protein sequences (16). PRATT returns patterns expressed in Prosite notation that include the sequence pattern, the number of sequences that match the pattern (“hits”) and a “fitness” score, which is a non-statistical parameter that reflects the non-randomness or complexity of the pattern. PRATT has several variable parameters that significantly influence the patterns found. One of the most important parameters is C% (or CM), the minimum number of unique sequences to match in a pattern. If C% is set too high, no patterns are found or only low fitness non-specific patterns are found. If C% is set too low, then many patterns are observed that are sub-patterns of a common parent pattern. As C% values decline, the length of the patterns and their fitness increases, depending on the specificity of the antibody. We illustrated the effect of C% values again using mOC1 (Table I). As the value of C% drops, the number of sequences satisfying the pattern (“hits”) drops and the fitness increases. The flexibility number (FN) specifies the maximum length or flexibility of arbitrary sequence spacers or “wild card” regions that can be any amino acid. We chose a FN value of 0 so that only fixed length patterns where X is one residue were evaluated. While PRATT returns the same patterns with larger values of FN, the program takes longer to run and PRATT also finds more low fitness patterns because of the increased flexibility. Pattern length (PL) is the maximum pattern length allowed and it must be set long enough to accommodate the longest epitope encountered. If it is set too low, PRATT will find multiple lower fitness sub-patterns that segments of a larger pattern. Longer values do not yield any more or different patterns, so we used a value of 12 because the maximum sequence length is 12. For most of the analysis we used C% = 10; FN = 0 and PL = 12. If using these parameters did not find any patterns with fitness values above those observed for the random sequence library alone (fitness > 12.50), then we adjusted the C% value down until higher fitness patterns were obtained.

**Table I.**
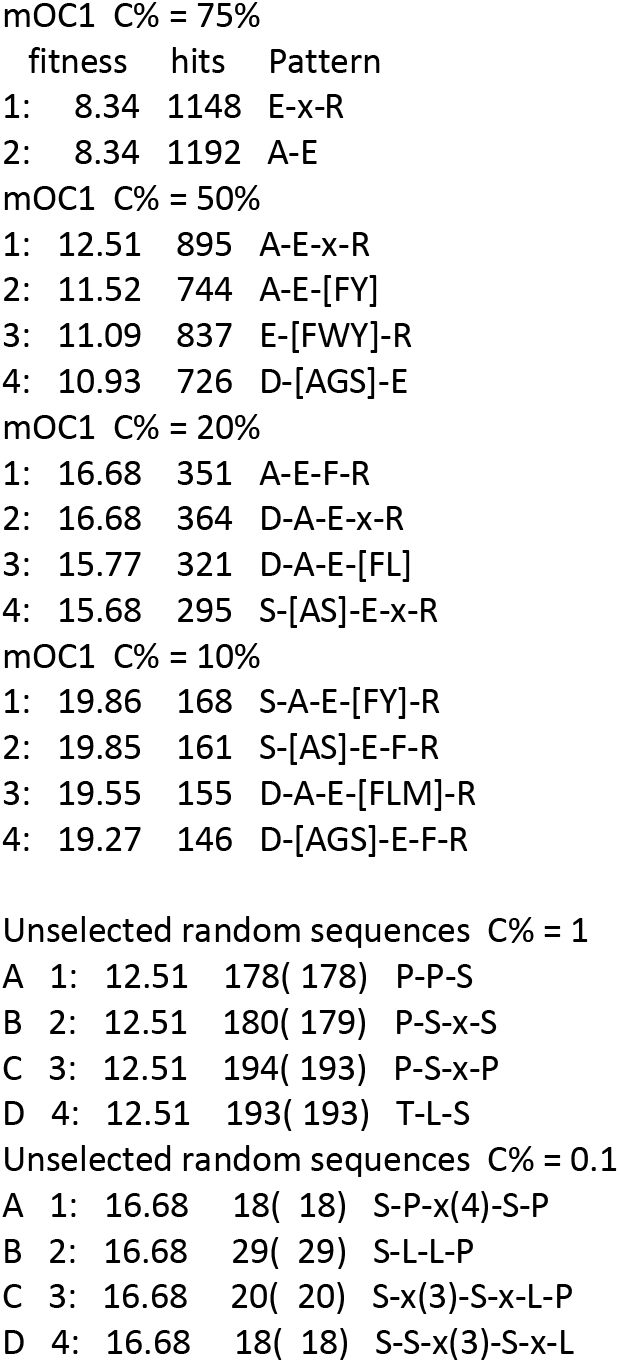
Effect of C% on patterns

|  | fitness | hits | Pattern |
| --- | --- | --- | --- |
| 1: | 8.34 | 1148 | E-x-R |
| 2: | 8.34 | 1192 | A-E |
| mOC1 C% = 50% |  |  |  |
| 1: | 12.51 | 895 | A-E-x-R |
| 2: | 11.52 | 744 | A-E-[FY] |
| 3: | 11.09 | 837 | E-[FWY]-R |
| 4: | 10.93 | 726 | D-[AGS]-E |
| mOC1 C% = 20% |  |  |  |
| 1: | 16.68 | 351 | A-E-F-R |
| 2: | 16.68 | 364 | D-A-E-x-R |
| 3: | 15.77 | 321 | D-A-E-[FL] |
| 4: | 15.68 | 295 | S-[AS]-E-x-R |
| mOC1 C% = 10% |  |  |  |
| 1: | 19.86 | 168 | S-A-E-[FY]-R |
| 2: | 19.85 | 161 | S-[AS]-E-F-R |
| 3: | 19.55 | 155 | D-A-E-[FLM]-R |
| 4: | 19.27 | 146 | D-[AGS]-E-F-R |
| Unselected random sequences C% = 1 |  |  |  |
| A 1: | 12.51 | 178( 178) | P-P-S |
| B 2: | 12.51 | 180( 179) | P-S-x-S |
| C 3: | 12.51 | 194( 193) | P-S-x-P |
| D 4: | 12.51 | 193( 193) | T-L-S |
| Unselected random sequences C% = 0.1 |  |  |  |
| A 1: | 16.68 | 18( 18) | S-P-x(4)-S-P |
| B 2: | 16.68 | 29( 29) | S-L-L-P |
| C 3: | 16.68 | 20( 20) | S-x(3)-S-x-L-P |
| D 4: | 16.68 | 18( 18) | S-S-x(3)-S-x-L |

Non-specific binding has a significant effect on C%, because the non-specific sequences do not fit a specific pattern, lowering the threshold number of C% required to observe the specific patterns. The non-specific sequences give the same low fitness patterns of 3-4 residues of the most abundant amino acids (S, L and P) that the unselected library does, with a fitness score of 12.5 for C% = 1 and 16.68 for C% = 0.1 (Table I). We first analyzed the effect of non-specific sequences by comparing the patterns found in the single copy sequences that contain the vast majority of non-specific sequences to the patterns obtained from the top 100 most frequently encountered sequences that bind in a sequence specific fashion. Surprisingly, there is very little difference in the patterns found, although the exact pattern and fitness score can vary slightly comparable to that caused by changing the C% value. Because there was little difference for all of the antibodies between the highly observed sequences and the total sequences, we show the data obtained from the total sequences here (Table II). Only the top 3 patterns with a fitness score above 12.5 are shown in Table II. If more than one pattern had the same fitness score, the patterns with the highest number of hits are shown.

**Table II.**
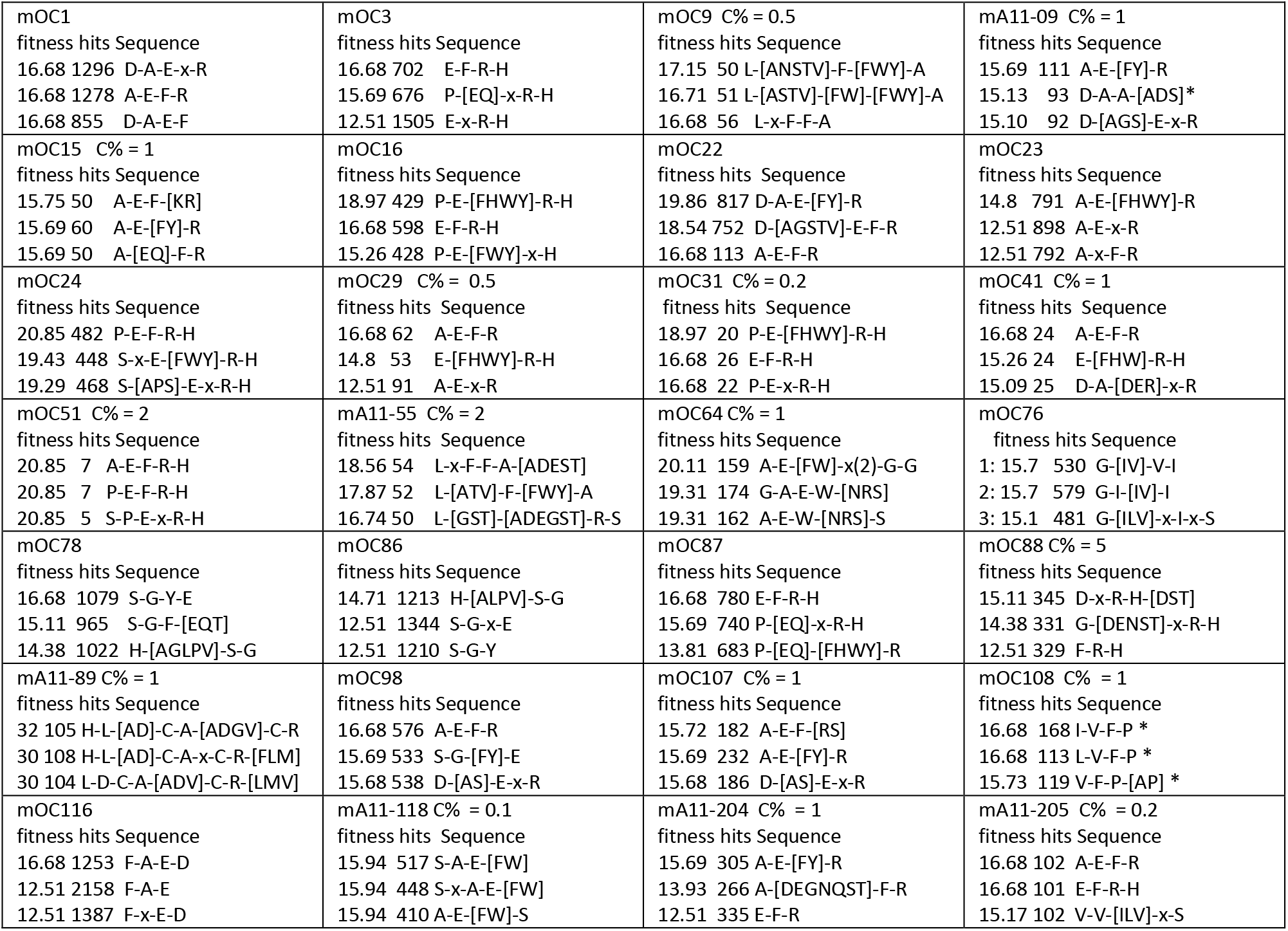
Major epitope patterns observed for 28 monoclonal antibodies. The 3 highest fitness Pratt patterns are shown. Positions denoted by x can be any amino acid. Positions denoted by square brackets denote the preferred amino acids at wild card positions. C% = 10 was used unless a value for C% is shown. * denotes patterns where a strong position effect is observed with the pattern beginning at the first residue of the sequence.

Half of the antibodies (mA11-09, mOC15, mOC29, mOC31, mOC41, mOC51, mA11-55, mA11-89, mOC98, mOC107, mOC108, mA11-118, mA11-204 and mA11-205) displayed no specific pattern at C% = 10, returning low fitness value pairs and triplets of amino acids that are similar in composition and fitness to the patterns observed for the unselected random library sequences that consist of the most abundant amino acids, such as leucine, proline, threonine and serine, indicating the patterns are dominated by non-specific sequences (Supplemental Information Table II). All of these antibodies display specific patterns at lower values of C% as listed in Table II. All of the mOC series antibodies and 5 of the 6 A11 antibodies (mA11-09, 55, 118, 204 and 205) display patterns related to the Aß sequence while mA11-89 displays a pattern unrelated to Aß. Individual peptides belonging to the mA11-89 patterns give multiple high identity hits in the non-redundant protein data base (data not shown). It is not clear why this antibody binds to Aß oligomers and oligomers from other amyloid sequences, although the sequence pattern observed may form a mimotope that is common to the ß-barrel oligomers that mA11-89 binds to.

Although most of the antibodies bind to the amino terminal third of Aß, each antibody displays differences in its binding preference in terms of the precise location on the target sequence, the identity of target amino acids required for binding, the location and composition of wild card positions where any amino acid or a subset of alternative residues are permitted and locations where non-target amino acids are preferred over target residues. Non-target amino acids such as a S or P at sequence position 1 or 2 are preferred or tolerated in several antibodies (S or P: mOC16, 24, 31, 51, 87 and mA11-118). Many of the antibodies that map to a region of Aß that contain an F or Y residue can accommodate either amino acid or sometimes W, while mOC116 only recognizes sequences containing the target F at position 20. Inspection of the actual sequences immunoselected by the antibodies indicate that mOC108 has a strong positional preference for sequences that begin at LVF or IVF, indicating that it prefers a free amino terminus. Sequences beginning at [IL]VF occur at approximately 93% of the total sequences recognized by mOC108.

Twenty-six of the monoclonals clearly recognize an Aß related epitope that is located in the amino-terminal (1–15) or central (16–30) thirds of the molecule or both (Fig. 2). Ten antibodies (mOC1, mA11-09, mOC15, mOC22, mOC23, mOC64, mOC98, mOC107, mA11-118 and mA11-204) all have epitopes centered on residues 1-5 DAEFR. Ten antibodies (mOC3, mOC16, mOC24, mOC29, mOC31, mOC41, mOC51, mOC87, mOC88 and mA11205) prefer residues 4-7 FRH, while mOC antibodies 78 and 86 bind to a site centered on residues 8-10 (SGY in Aß). Four monoclonal antibodies bind to epitopes in the central third of Aß. mOC108 binds to 17-19 (LVF), mOC9 and mA11-55 prefer the residues LVFFA (residues 17-21) and mOC 116 binds to residues 20-23 (FAED). One antibody, mOC76 gives a pattern, G-[IV]-[IV]-I-x-S, that is related to carboxyl terminal sequence residues 38-43 of Aß42, even though our previous epitope mapping study indicated that this antibody maps to residues 5-10, RHDSGY using the “PepSpots” overlapping 10mer epitope mapping method (10).

**Figure 2.**
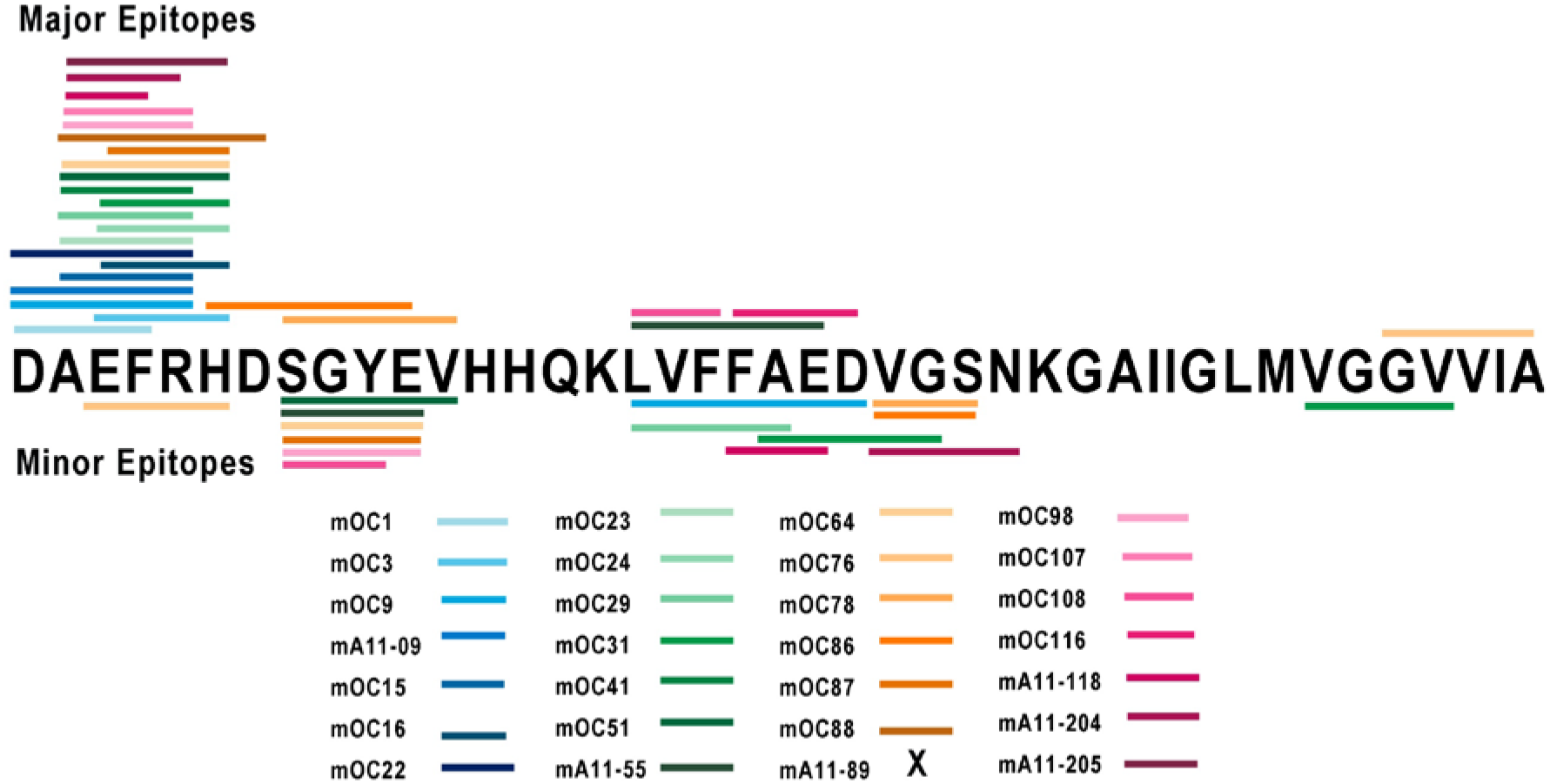
Map of antibody binding sites on Aß42. The binding sites identified in Tables II and III are shown as color coded bars. The major epitope is shown by bars above the Aß sequence and the minor epitope for antibodies that bind to more than one Aß segment is shown by the bars below the Aß sequence. No binding site is indicated for mA11-89 because it does not bind to a linear Aß sequence by epitomic analysis.

**Table III.**
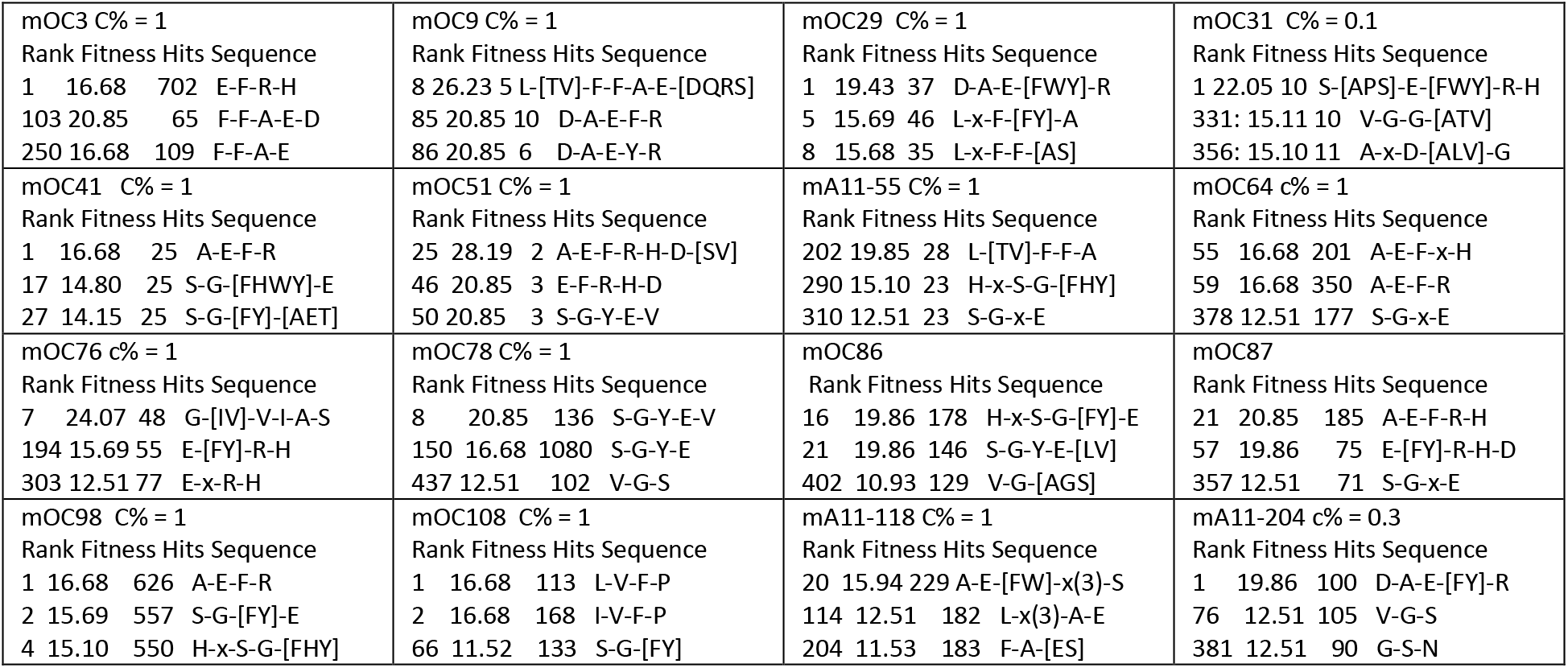
Antibodies that recognize more than 1 non-overlapping segment of Aß

Our previous studies indicated that 6 of the antibodies (mOC1, 24, 31, 51, 78 and 98) recognize a discontinuous epitope because they react with more than one segment of the Aß sequence using the overlapping peptide PepSpots method (17). The epitomic analysis of the antibodies also provides support for the recognition of discontinuous epitopes consisting of non-overlapping sequence segments of Aß. For example, antibody mOC98 binds to both residues 2-5 and residues 8-11 as the top two patterns observed (Table II). In consideration of the possibility that other Aß segments of discontinuous epitopes may be lower ranked patterns, we ran the PRATT analysis at lower values of C% (1%) and output all of the patterns with a fitness score above 12.5. We then searched for patterns containing other segments of Aß. This analysis identified 16 antibodies that recognize more than 1 non-overlapping segment of Aß as shown in Table III. While the epitomic analysis confirms the discontinuous epitopes identified by the PepSpots assay for antibodies mOC3, 51, 78 and 98, it fails to confirm a second binding segment for mOC1, and 24. In addition, it identifies 12 additional antibodies with discontinuous epitopes that were not observed using the PepSpots membranes (mOC9, 29, 31, 41, 64, 76, 86, 87, 98, and mA11-55, 118 and 204). The resulting map of antibody binding sites on Aß including the major epitopes for 27 of the 28 antibodies and the minor epitopes for the antibodies that recognize more than 1 segment of Aß is shown in Figure 2. Only two segments of the Aß sequence are not recognized by any of the antibodies, residues 13-16 (HHQK) and residues 28-35 (KGAIIGLM).

In order to confirm the binding sites for the antibodies on the Aß target sequence and attempt to resolve the conflicts between the epitomic approach and the PepSpots method, we repeated the peptide array using synthesized overlapping 10mers of the Aß sequence from the −4 position to residue 45 and arraying the sequences on glass slides and probing the array with the antibodies. The data is shown in Figure 3 and a comparison of results of the three different methods is summarized in Table IV. The microarray data indicates that only 5 of the 16 antibodies identified by epitomic analysis as binding to discontinuous epitopes are confirmed by the two peptide array assays (mOC31, mOC51, mOC78, mOC98 and mA11-55). Even though all 3 approaches agree, the actual segments are different among the three different assays for some of the antibodies. For example, mOC51 binds to residues 3-7 and 19-25 according to PepSpots, residues 6-11 and 17-21 according to Epitomic analysis and 13-21 and 28-37 by the microarray data. The two peptide arrays don’t agree with each other for several of the antibodies. The epitomic analysis finds more discontinuous epitopes than the peptide array approach and this may be due to the fact that in random sequence selection, both segments may be present simultaneously to create a stronger binding pair than the individual segments by themselves. The microarray results also confirm that mOC76 binds to the carboxyl-terminal sequences of Aß, binding most strongly to the peptide ending at residue 45 (GVVIATVI) and fails to confirm the data from the PepSpots assay that indicate binding at the amino terminus.

**Table IV.** Summary of discontinuous epitopes

| Table IV Summary of discontinuous epitopes |  |  |  |
| --- | --- | --- | --- |
|  | PepSpots | Epitomics | Array |
| mOC1 | + | - | + |
| mOC3 | - | + | + |
| mOC9 | - | - | - |
| mA11-09 | - | + | - |
| mOC15 | - | - | - |
| mOC16 | - | - | - |
| mOC22 | - | - | - |
| mOC23 | - | - | - |
| mOC24 | + | - | - |
| mOC29 | - | + | - |
| mOC31 | + | + | + |
| mOC41 | - | + | - |
| mOC51 | + | + | + |
| mA11-55 | + | + | + |
| mOC64 | - | + | - |
| mOC76 | - | + | - |
| mOC78 | + | + | + |
| mOC86 | - | + | - |
| mOC87 | - | + | - |
| mOC88 | - | - | + |
| mA11-89 | - | - | - |
| mOC98 | + | + | + |
| mOC107 | - | - | - |
| mOC108 | - | + | - |
| mOC116 | - | - | + |
| mA11-118 | - | + | - |
| mA11-204 | - | + | - |
| mA11-205 | - | - | - |
| + indicates a discontinuous epitope |  |  |  |

**Figure 3.**
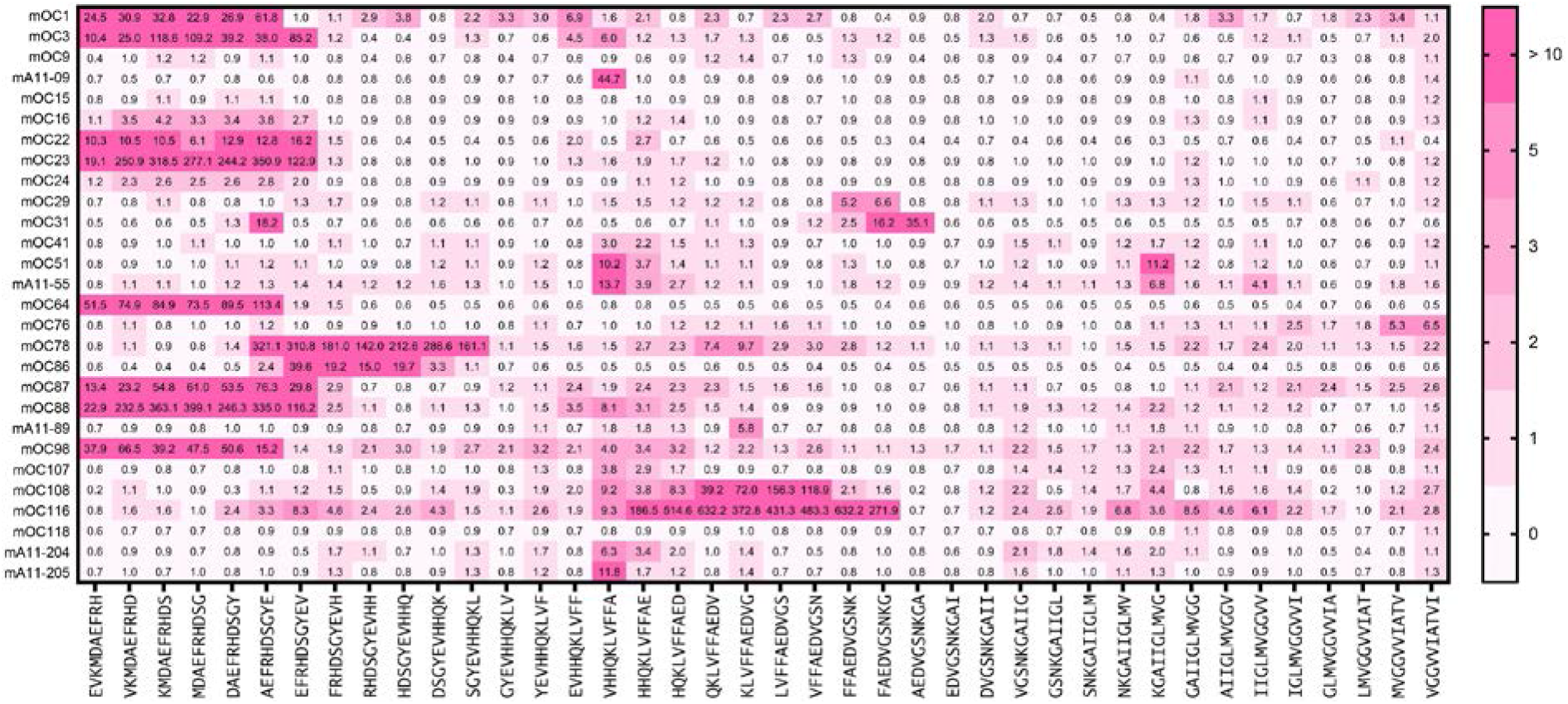
Microarray analysis of overlapping Aß peptide 10mer sequences from −4 to 45 as indicated along the bottom of the figure. The spot intensity is indicated by pink to red boxes containing the fold increase above background of the binding.

Most of the antibodies are specific for aggregated forms of Aß and do not recognize monomer or the amyloid precursor protein, so it is not obvious from the epitope fine structures why the sequences displayed by the phage bind. It is possible that the monomeric sequence binds weakly in comparison to the aggregated form or that the random sequences fused to pIII aggregate on the phage. To investigate this issue further, we analyzed the specific sequences immunoselected for the propensity to form amyloid aggregates by three different algorithms, AGGRESCAN (18), Waltz (19) and the 3D profile method (20). The first two methods are sequence-based while the last method is structure based. The three methods give a general agreement on the amyloid forming propensity, but the Waltz algorithm provides higher estimates and the 3D profile methods gives lower estimates of amyloid formation. We analyzed the top 100 sequences for each antibody and 100 unselected random sequences for amyloid formation propensity and the results are shown in Table V. Although there is considerable variation in the amyloid-forming propensity for some of the sequences, the antibodies that are specific for amyloid aggregates are associated with sequences that have a significantly higher propensity for forming amyloid than the antibodies that recognize monomer (p < 0.002). Since there are about 5 copies of the pIII protein all located on the head side of the filamentous phage, it is conceivable that they could form amyloid-like intermolecular aggregates on the surface of the phage. Some of the antibodies that recognize monomer have sequences with very low amyloid forming propensity that is lower than the average amyloid forming propensity for random 12mers from the unselected library and lower than that for the proteins, APP, tau, α-synuclein and BSA. This preference for low amyloid forming propensity may be a reflection of the preference of these antibodies for unstructured regions of peptide. We have previously reported that some of these antibodies bind to amyloid plaques and intraneuronal amyloid in human and transgenic mouse brain while others do not (10,21). In addition, mOC31 specifically recognizes vascular amyloid (10,22). We also found that two of the antibodies, mOC22 and mOC23 differentially recognize distinct regions of cored plaques (Fig. 4). mOC22 recognizes the entire cored plaque while mOC23 preferentially binds to the rim of the plaque, indicating that even though both antibodies bind to the same amino terminal region of Aß, their distinct epitomic specificities are meaningful in their ability to identify inhomogeneities in plaque structure. Diffuse plaques stain uniformly with both antibodies indicating that different types of plaques display different sets of epitopes (Fig. 4). The staining properties of antibodies mA11-09 and mA11-89 in human brain have not yet been reported. We found that mA11-09 stains intraneuronal amyloid that co-localizes with 6E10, while neither antibody stains plaques (Fig. 5), consistent with the reported properties of A11 serum (23), and provides further evidence for the utility of these antibodies in distinguishing novel types of amyloid in human brain.

**Table V.** Aggregation propensity of immunoselected peptides.

| Name | Waltz<br>% hits | AggScan<br>% hits | 3D %<br>hits | Average |
| --- | --- | --- | --- | --- |
| RandomSeq | 32 | 28 |  |  |
| BSA | 41.5 | 31.6 |  |  |
| APP771 | 35.8 | 26.5 |  |  |
| Synuclein | 25.7 | 51.4 |  |  |
| mOC1 | 3 | 12 | 11 | 8.7 |
| mOC3 | 29 | 13 | 5 | 15.7 |
| mA11-09 | 52 | 36 | 15 | 34.3 |
| mOC9 | 63 | 50 | 23 | 45.3 |
| mOC15 | 42 | 36 | 15 | 31.0 |
| mOC16 | 23 | 2 | 7 | 10.7 |
| mOC22 | 33 | 20 | 11 | 21.3 |
| mOC23 | 25 | 13 | 3 | 13.7 |
| mOC24 | 16 | 5 | 3 | 8.0 |
| mOC29 | 58 | 47 | 28 | 44.3 |
| mOC31 | 38 | 29 | 19 | 28.7 |
| mOC41 | 54 | 47 | 32 | 44.3 |
| mOC51 | 51 | 32 | 9 | 30.7 |
| mA11-55 | 46 | 34 | 23 | 34.3 |
| mOC64 | 58 | 22 | 11 | 30.3 |
| mOC76 | 69 | 80 | 58 | 69.0 |
| mOC78 | 54 | 34 | 25 | 37.7 |
| mOC86 | 51 | 44 | 29 | 41.3 |
| mOC87 | 25 | 5 | 5 | 11.7 |
| mOC88 | 20 | 9 | 4 | 11.0 |
| mA11-89 | 66 | 72 | 42 | 60.0 |
| mOC98 | 22 | 13 | 9 | 14.7 |
| mOC107 | 44 | 40 | 17 | 33.7 |
| mOC108 | 51 | 40 | 9 | 33.3 |
| mOC116 | 65 | 43 | 9 | 39.0 |
| mA11-118 | 36 | 33 | 16 | 28.3 |
| mA11-204 | 46 | 36 | 16 | 32.7 |
| mA11-205 | 51 | 43 | 20 | 38.0 |
| 4G8 | 53 | 79 | 21 | 51.0 |
| 6E10 | 23 | 7 | 4 | 11.3 |

**Figure 4.**
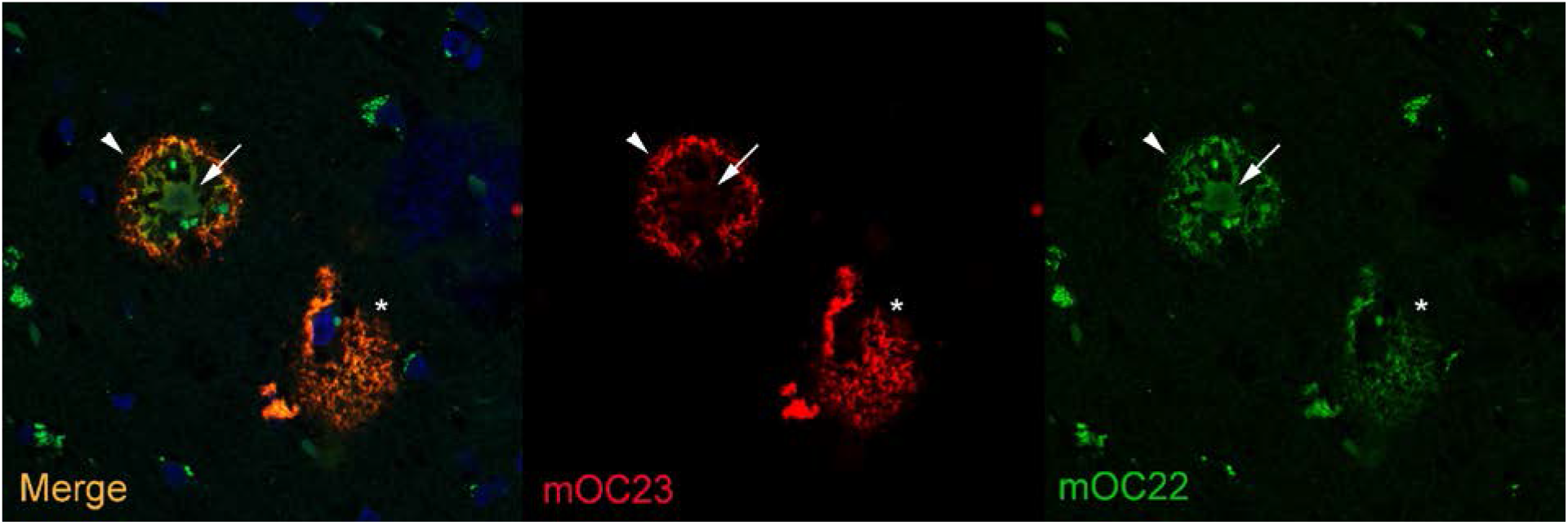
Differential staining of plaques in human AD brain by mOC22 (green) and mOC23 (red). mOC22 stains the entire cored plaque (arrow) while mOC23 only stains the rim (arrow head). A diffuse plaque is stained by both antibodies (Asterix). The cored plaque is approximately 50 micrometers in diameter.

**Figure 5.**
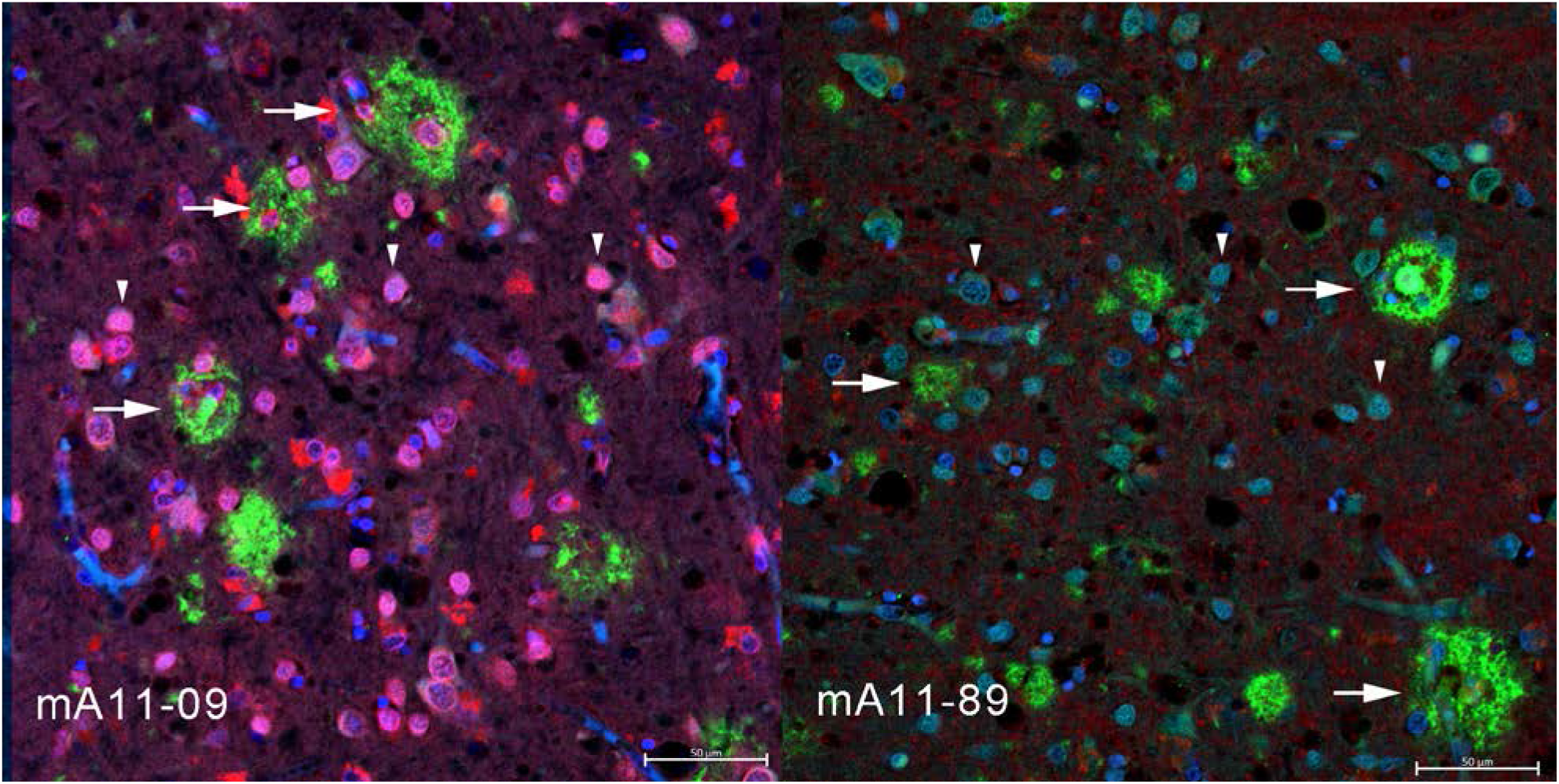
Monoclonal antibody mA11-09 stains intraneuronal amyloid and both mA11 and mA11-89 fail to stain amyloid plaques. The intraneuronal mA11-09 immunoreactivity (arrow heads) largely overlaps with 6E10 Aß immunoreactivity (left panel), which does not stain with mA11-89 (right panel). Neither antibody stains amyloid plaques that are stained by 6E10 (arrows). Scale bar = 50 um.

## Discussion

The immune response to amyloid Aß is highly diverse, giving rise to many antibodies that bind to the same regions of the Aß sequence and yet differentially react with specific aggregation states and structures in vitro and identify distinct types of amyloid deposited in human and transgenic mouse brain. We hypothesized that this diversity in immunoreactivity may reflect the structural diversity that the Aß peptide can adopt upon aggregation into ß-sheet amyloid and that differences in the way in which the antibody binds to the different structures would be reflected in the pattern of amino acids they prefer for binding. We found that the antibodies display distinct differences in terms of the location and identity of the target sequence residues that are critical for binding, the non-overlapping segments that participate in binding and in the preference for non-target residues. The results show that all of the antibodies display distinct preferences for epitopes consisting of a mixture of target and non-target residues and positions where a number of different amino acids are accommodated while other amino acids are rejected. This suggests that the regions of Aß containing these epitopes are able to adopt a number of different structures such that the epitopes are displayed or not in a combinatorial fashion. This provides an explanation for why the antibodies bind to different Aß fibril structures (13) and bind to different types of amyloids in vitro and in vivo (10,22,23,26). Amyloids are intermolecularly hydrogen bonded cross-ß structures that are structurally polymorphic (reviewed in (27,28)), and the fact that Aß displays a number of different structural polymorphisms is consistent with the fact that the immune response gives rise to a large number of different antibodies that have different binding modes for the different structures of the same peptide segments of Aß. The fact that so many of the antibodies that bind to the amino terminal segment of Aß are specific for aggregated Aß and recognize different structural variants indicates that the commonly held view that antibodies that bind to a linear segment of a peptide sequence are “non-conformational” is an over simplification at least for anti-amyloid antibodies. A more accurate view may be whether it can adopt more than one structure. Mimotopes such as the ones identified here may give a more predictable immune response because it may adopt a single predominant structure and give rise to a more predictable immune response.

Additionally, we found that the antibodies differ significantly in the non-overlapping segments of the Aß peptide that constitute the epitope that the antibody preferentially binds to. Most of these discontinuous segments are located in the amino terminal or central thirds of the sequence and suggest a combinatorial mechanism for the specificity of the antibodies that bind to discontinuous epitopes where different segments combine to form the epitope. A standard approach for epitope mapping is the PepSpots assay, where a series of overlapping peptides are synthesized and arrayed and used to determine the region of protein sequence to which an antibody binds. While this is a reasonably inexpensive and facile method, it lacks fine details of the interactions and fails to identify many of the discontinuous conformational epitopes revealed by the epitomic approach, such as those that are found on amyloid aggregates. Phage display has also been used to identify epitopes, but the classical approach of sequencing positive individual clones limits the number of samples one can realistically process at once (24). Here we combine random sequence phage display and next generation sequencing to identify the fine details of monoclonal antibody recognition of Aß and its aggregates. Thousands of immunoselected sequences were analyzed using the PRATT 2.1 pattern recognition algorithm for patterns common to specifically bound phage that define the preferred epitope sequences recognized by the antibody. By coupling immunoselection of random sequences with deep sequencing, the results provide unprecedented insight into the nature of the epitope and its interaction with the antibody. Not only do the results identify the residues of the target sequence that constitute the binding site, but they also provide details about non-target residues that are allowed, disallowed or preferred for binding. Epitomic analysis also identifies non-overlapping segments that may constitute conformational epitopes for antibodies that fail to bind to these sequences using peptide array methods. This detailed structural information about the epitope may be valuable for modeling the atomic details of the antibody-epitope interaction.

Identifying conformational epitopes for monoclonal antibodies is an important and challenging topic for immunology and epitomic analysis may be a rapid and facile means of investigating conformational specificity. The availability of multiple peptide sequences simultaneously for antibody binding may allow the identification of binding segments where each individual sequence binds too weakly to allow identification using single linear peptide segment approaches, like peptide arrays. In addition, some antibodies appear to bind one segment more tightly that the other so they may be misidentified as having a single linear epitope by peptide arrays when they actually recognize a discontinuous epitope. Reexamination of these antibodies using epitomic analysis may be warranted. These details may be very useful for understanding the binding mechanism and modeling the binding interaction in the absence of detailed structural information of the antibody-antigen complexes. This insight could also be very useful for predicting antibody cross reactivity.

We hoped to also gain some insight into some of the peculiar properties of anti-amyloid antibodies, such as why many of them are specific for aggregates, why they recognize generic epitopes that do not depend on a specific amino acid sequence and why the A11 and OC series antibodies recognize mutually exclusive aggregation-specific epitopes. A11 binds known repeating antiparallel β structures, such as antiparallel Aß42 prefibrillar oligomers (25), hemolysin pores (15) and β cylindrins (26). Antibodies in A11 serum also bind to α-sheet structures (27). OC binds to known parallel, in-register structures, such as Aß (13), α-synuclein and IAPP fibrils (8). In antiparallel ß sheets the strands alternate in opposite orientations, so it seems more likely that antibodies that are specific for antiparallel ß sheets would bind to discontinuous epitopes. Indeed, half of the A11 antibodies appear to bind to more than one Aß segment and the combinatorial nature of the discontinuous segments provides an additional layer of specificity to the antibodies with these epitopes. However, approximately half of the mOC series monoclonals also display discontinuous epitopes, so this feature alone does not explain specificity of the A11 and OC antibodies. If the OC antibodies bind to parallel ß sheets, then it would seem to indicate that these antibodies must bind to the ends of the beta sheet hair pin which is the only part of the fibril structure where both segments would be exposed for antibody binding.

Monoclonal antibodies against Aß are an important class of therapeutic agent under development for AD. Although many antibodies have been tested in the clinic, only two, Aducanumab and BAN2401, have been reported to slow the progression of the disease, but they have yet to demonstrate consistent therapeutic activity and have not yet been approved for human use (28–30). It is not yet clear what distinguishes these antibodies from the ones that did not demonstrate clinical effectiveness, but they are claimed to specifically target aggregated forms of Aß. We have previously reported that the majority of the antibodies examined here are specific for aggregation-related epitopes and do not bind Aß monomer or APP as determined by Western blotting and dot blots on preparations of synthetic Aß oligomers, fibrils and monomer (9,10). While these approaches are facile, they may not reproduce the structures that exist in vivo and they are complicated by the difficulty of preparing homogeneous populations of aggregates. Analysis of the immunoselected sequences by amyloid prediction algorithms provides an independent and unbiased means of assessing the specificity for aggregates. The data indicate that the sequences that bind to aggregation-specific antibodies have a significantly higher predicted propensity to aggregate than the random sequences as a whole or the sequences preferred by the antibodies that bind to Aß monomer. This suggests that the random sequence displayed by the phage pIII protein may be aggregated even though there are a relatively low number of copies of the pIII protein. It is interesting that many of the antibodies that bind monomeric Aß show a lower than average propensity to aggregate. It is not yet clear whether this means that they are more likely to be unstructured, which facilitates the induced fit in the antigen combining site or whether they adopt a common non-aggregating structure.

The information obtained by epitomic analysis can also facilitate the investigation of which specific antibodies against Aß occur in human serum and whether they are associated with avoiding or predictive of having AD. We found that the antibodies display a unique pattern of amino acid residues that is required for binding, positions where any amino acid or a restricted set of amino acids are permitted and residues that are incompatible with binding. This pattern is unique to the antibody and is most likely determined by the structure of the complementary paratope. These unique patterns could be very useful to serve as a fingerprint to identify similar antibodies that recognize the same epitope in complex mixtures of antibodies, such as in human blood or CSF. It may be possible to synthesize a set of peptide epitopes and mimotopes unique to the individual antibodies and screen human serum for immunoreactivity at these sequences. This would allow the identification of antibodies that are correlated with protection against AD or serve as biomarkers of AD. It could also enable the identification of disease subtypes and precision medicine approaches to AD immunotherapy that target the specific polymorphic structures that occur in human brain. With the large number of different antibodies with unique epitopes and specificities for polymorphic amyloid structures, there are many antibodies that remain to be examined in human clinical trials.

## Experimental procedures

### Antibodies

Twenty eight rabbit monoclonal antibodies consisting of 5 antibodies derived from A11 serum and 23 derived from OC serum were produced and characterized as previously described (9,10). Eight antibodies are commercially available from Abcam mOC22 (cat# ab205339), mOC23 (cat# ab205340), mOC31 (cat# ab201059), mOC64 (cat# ab201060), mOC78 (cat# ab205341), mOC87 (cat# ab201062), mOC98 (cat# ab201061) and mOC116 (cat# ab205342). All of the other antibodies are available under material transfer agreement upon request.

### Phage display

Phage display was performed using New England Biolab’s Ph.D.™-12 Phage Display Peptide Library Kit (Cat# E8110S) following manufacturer’s instructions. Briefly, 1×10^11^ CFU where incubated with 1 μg of purified monoclonal antibody in a final volume of 200 μl of TBS-T (TBS with 0.5% tween-20) for 20 minutes at room temperature. After the incubation period, the sample was resuspended in 50 μl of Protein A-coated magnetic beads (Novex DYNAL Dynabeads Protein A, cat# 10002D). Following a 20-minute incubation, the unbound phage was removed, and the beads washed with 1 ml of TBS-T. The beads were then resuspended in 200 μl of TBS-T, placed in a 96 well plate and washed in TBS-T using a Thermofisher Kingfisher magnetic particle processor. Bound phage was eluted in 200 μl glycine buffer pH 2.2 and immediately neutralized by adding 20 μl of 1 M Tris pH 9. One hundred μl is saved to isolate phage DNA (unamplified panning) and the rest is used to infect 1.5 ml of LB broth with *E coli* and amplified at 37°C. After 4.5 hours, the bacterial broth is centrifuged for 10 minutes at 18k x *g* in a microcentrifuge, the supernatant is recovered and the phage is precipitated overnight at 4°C by adding 250 μl of 20% PEG 8000/2.5M NaCl. The following day, the samples are centrifuged 10 minutes at 18k x *g* at 4°C and the phage pellet is resuspended in 100 μl of TBS. This is the amplified panning, and 10 μl are used for antibody selection in the next panning.

### Phage DNA isolation and library preparation

Phage DNA was isolated using a standard phenol:chloroform method (31). Quality was assessed by visualization in a 1% agarose gel, and its concentration measured by spectrophotometry. One hundred ng of phage DNA were used as template for PCR amplification for the Next Generation Sequencing step. The phage DNA amplicons were barcoded, pooled and a 10 nM library was sequenced commercially on an Illumina MiSeq platform. (Laragen Inc, Culver City, California, USA).

### Data analysis

The Illumina sequencing data was processed by a BASH script (Supplemental Information) that extracts the DNA sequence coding for the dodecapeptides, translates them to protein, counts how many times each unique sequence was found in the sequencing file (frequency) and removes common background sequences due to unspecific binding to the protein A beads. The peptide sequences were sorted by frequency and converted to FASTA format as previously described (32). The FASTA sequences were written with an identifier label that contains the antibody name, the unique sequence number and the number of times the sequence was observed in the following pattern: >(antibody)*(sequence #)_(frequency) to facilitate machine counting of the frequency. The sequences were analyzed to determine the amino acid sequence patterns they contain using PRATT 2.1 (16) that was edited and recompiled to accommodate up to 200,000 sequences as previously described (32).

### Microarray analysis

For microarray fabrication, biotinylated amyloid peptides spanning the amyloid beta sequence (−5 to 45) were pre-incubated with NeutrAvidin (ThermoFisher Scientific, MA, USA) in phosphate-buffered saline (PBS) at molar ratio of 4:1 for 1 hr at room temperature with gentle agitation. Final concentration of peptides in the complex was 0.5 mg/ml. Tween 20 (T-PBS) and glycerol was added to the peptide complex solution at final concentration of 0.001%, and then the solution was spotted onto nitrocellulose coated glass AVID slides (Grace Bio-Labs, Inc., OR, USA) using an Omni Grid 100 contact microarray printer (Genomic Solutions). One nanoliter of peptide solution was delivered onto the membrane, corresponding approximately 0.5 ng of peptide per spot. Slides were stored in desiccator until use. Monoclonal antibodies were diluted in protein array blocking buffer (GVS, Sanford, ME, USA) to a final concentration of 5 ng/ml. Concurrently, arrays were rehydrated in blocking buffer for 30 min. Blocking buffer was removed, and arrays were probed with samples using sealed chambers to avoid cross-contamination between the pads. Arrays were incubated overnight at 4°C with gentle agitation. Arrays were washed three times with washing buffer, Tris-buffered saline (TBS) containing 0.05% Tween 20 (T-TBS), and bound antibodies were detected by Cy5-conjugated goat anti-rabbit IgG (Jackson ImmunoResearch Laboratories, Inc., West Grove, PA, USA), diluted 1:200 in blocking buffer. After 1h at room temperature (RT) incubation, arrays were then washed three times with T-TBS and once with water. They were air dried by centrifugation at 500 × *g* for 10 min. Images were acquired using Perkin Elmer ScanArray Express HT confocal laser scanner at a wavelength of 670 nm and signal intensities were quantified using ProScanArray Express software (Perkin Elmer, Waltham, MA). All signal intensities were corrected for spot-specific background.

### Fluorescence microscopy

Human brains obtained from UCI Alzheimer’s Disease Research Center (UCI ADRC) Tissue Repository were sliced into 50 μm thick slices using a Vibratome Leica 1000 and stored in 1X phosphate-buffered saline (PBS) with 0.02% NaN3 (v/v). Antibody mOC22 was biotinylated with NHS-biotin (ThermoFisher EZ-Link Catalog # 20217) according to the manufacturer’s instructions. For antibodies mOC22 and mOC23, the sections were incubated overnight at room temperature in at a concentration of 0.1 μg/ml in TBS containing 1% BSA and then washed 3 times in TBS. The sections were incubated with goat anti rabbit IgG labeled with AlexaFluor 647 (A32728, Thermo Fisher Scientific), 1 μg/ml in TBS with 1% goat serum for 1 hr. Sections were washed 3 times in TBS for 5 minutes and incubated in TBS containing 1% rabbit serum for 30 minutes. The sections were then incubated overnight in 0.1 μg/ml biotinylated mOC22 and then washed 3 times in TBS, incubated in TBS-1% BSA and incubated with 1 μg/ml AlexaFluor 488 streptavidin ThermoFisher S21374)for 1 hr, then washed 3 times with TBS and mounted on a slide and imaged with a Zeiss LSM 700 laser confocal microscope. For antibodies mA11-09 and mA11-89, sections were subjected to antigen retrieval with 1M citric acid (pH 6.0), heated by microwave for 2 min, followed by 4 min incubation with 80% formic acid. Sections were washed 3 times with PHEM buffer (60mM Pipes, 25 mM Hepes, 10mM EGTA, 4 mM MgSO4), followed by permeabilization with PHEM with 0.2% TritonX-100 (v/v) for 20 min, shaking at RT. Afterward, sections were blocked in PHEM with 2% BSA and 1.2% normal goat serum for 1 h at RT, shaking. Amyloid was stained overnight at RT, shaking with (1:100) hybridoma supernatant from monoclonal antibodies in the blocking solution. The omission of the primary antibodies was taken along as negative controls. Further, brain sections were subjected to a series of washes with PHEM buffer and incubated with the secondary antibody Alexa Fluor-647 (A21245, Thermo Fisher Scientific) (1:300) in PHEM buffer with 2% BSA for 3 h at RT, shaking in the dark. Next, the sections were washed with PHEM buffer and stained with 5 μg/mL DAPI solution for 10 min at RT. Stained brain sections were then mounted using ProLong Diamond Antifade mounting medium (P36970, Thermo Fisher Scientific) and scanned with Zeiss LSM 700 laser confocal microscope.

## Data Availability Statement

All data files can be accessed at: Dryad, Dataset, https://doi.org/10.7280/D1QH5W

## Acknowledgments

Authors wish to thank Dr. Jeffrey Glabe for help with modifying and recompiling PRATT, Mr. Matt Gargus, Mr. Noe Neira Garza and Ms. Dania Alvarez for helping to write the BASH script to extract the peptide sequences from the sequencing files.

## Funding and additional information

This work was supported by NIH grants AG056507 and AG061857 (CG), and the Cure Alzheimer Fund (CG).

## Conflict of interest

The authors declare no conflicts of interest.

## Notes

### Competing Interest Statement

The authors have declared no competing interest.

### Summary of Updates

Revised introduction and discussion.

