## Supplemental Information for "Epitomic analysis of the specificity of conformation-dependent, anti-Aß amyloid monoclonal antibodies" (from another version of this preprint: version 2, 2020-10-28)

Supplemental Information Table I Nucleotide Frequency

| Nucleotide | Percent |
|---|---|
| G | 26.39 |
| A | 18.15 |
| T | 34.24 |
| C | 21.22 |

Supplemental Information Table II Amino Acid Frequency

| Amino Acid | Frequency | Amino Acid | Frequency |
|---|---|---|---|
| S | .109 | D | .039 |
| P | .103 | Y | .038 |
| L | .092 | Q | .036 |
| T | .092 | I | .035 |
| A | .069 | F | .031 |
| H | .052 | M | .029 |
| R | .052 | E | .027 |
| V | .050 | K | .025 |
| N | .048 | W | .019 |
| G | .042 | C | .010 |

Bash script for processing sequencing files.

```
#unzips any file with a .gz at end
for file in /data/users/UserName/UserDirectory/job_trials/*.gz; do gunzip ${i};done

#Name A11
for file in 1_* 2_* 3_*; do mv "$file" "${file/S*_/A11.fastq}"; done

#Name OC
for file in 21_* 24_* 14_*; do mv "$file" "${file/S*_/OC.fastq}"; done

#Name Lock
for file in 15_* 20_*; do mv "$file" "${file/S*_/Lock.fastq}"; done

#name group Beads
for file in 96_* 97_* 98_*; do mv "$file" "${file/S*_/Bead.fastq}"; done

#Prinseq execute in Perl
for file in *.fastq; do prinseq-lite.pl -fastq ${file} -out_format 3;done

#removes all files EXCEPT files that have good in name from prinseq output
find . -type f ! -name "*good*" -exec rm -rf {} \;

#search all files that have good and ends with .fastq for TTCTCACTCT and next 36 basepairs
for file in *good*.fastq; do grep -o -P 'TTCTCACTCT.{0,36}' ${file}> ${file}_hit.txt; done

#removes all files EXCEPT .txt files in directory
find . -type f ! -name "*.txt" -exec rm -rf {} \;

#searches files that end with _hit.txt and removes TTCTCACTCT wherever its found
for file in *_hit.txt; do sed "s/TTCTCACTCT//g" ${file} > ${file}_final_thirtysix.txt; done

#trims name of files with prinseq_good and ends with .txt and replaces and renames anything
after prinseq_good as dna_sequence
for file in *prinseq_good*.txt; do mv "$file" "${file/prinseq_good*/dna_sequence}"; done

#for files in specified directory that end with dna_sequence run python script translate.py and
name the output the same with _translated added
for file in /data/users/UserName/UserDirectory/job_trials/*dna_sequence; do python
/data/users/UserName/UserDirectory/translate.py ${file}>${file}_translated; done

#removes all files EXCEPT ones with translated at the end
find . -type f ! -name "*translated" -exec rm -rf {} \;
```

```
#
find /data/users/UserName/UserDirectory/job_trials -type f -exec sed -i 's/*/X/g' {} \;

#
find /data/users/UserName/UserDirectory/job_trials/ -type f -exec sed -i '/.\{12\}/!d' {} \;

#THIS NEEDS ATTENTION! Bead background files need to be designated input!!!
for file in *Bead* ; do cat "$file" >> Bead_background.txt && rm "$file" || break ; done

#Remove/replaces lines that appear in bead background file and removes from files with
translated in name
for file in *translated ; do grep -vxFf *background.txt ${file}> ${file}_bead_filter.txt; done

#These lines provide individual files ranked sorted counted and labeled
for i in *bead_filter.txt; do
cat ${i} | sort | uniq -c | sort -nr >> ${i}_bead_ranked.txt;
done
for i in *dna_sequence_translated_bead_filter.txt_bead_ranked.txt; do
g=$(basename $i dna_sequence_translated_bead_filter.txt_bead_ranked.txt);
sed -e "s/[0-9]*//g" ${i} | sed -e "s/ //g" > ${g}_peptides.txt
awk -v g="$g" '{ print ">"g"_peptide_"NR"_"$1$0}' ${i} | awk '{print $1}' > ${g}_headers.txt;
paste -d '\n' ${g}_headers.txt ${g}_peptides.txt > ${g}_complete.txt
rm ${g}_peptides.txt
rm ${g}_headers.txt
done

#concatenates all A11 files
cat *A11* > A11_all.txt

#concatenates all OC files
cat *OC* > OC_all.txt

#concatenates all Lock files
cat *Lock* > Lock_all.txt

#aruably the most rigorous lines. This section takes the all.txt files creates a low counts file for
peptides that appear less than 5 and then also make a file for lines in each file that appear 5 or
more time. This then creates some temporary files to create a .fasta file with the peptides labeled,
ranked, with count in the header.
for i in *all.txt; do
cat ${i} | sort | uniq -c | sort -nr >> ${i}_sorted_ranked.txt;
done
for i in *_sorted_ranked.txt; do
awk '($1 > 4 ) ' ${i}>${i}_five.txt;
done
```

```
for i in *_sorted_ranked.txt; do
awk '($1 < 5 ) ' ${i}>${i}_low_counts.txt
done
for i in *_all.txt_sorted_ranked.txt_five.txt; do
g=$(basename $i _all.txt_sorted_ranked.txt_five.txt);
sed -e "s/[0-9]*//g" ${i} | sed -e "s/ //g" > ${g}_peptides.txt
awk -v g="$g" '{ print ">"g"_peptide_"NR"_"$1$0}' ${i} | awk '{print $1}' > ${g}_headers.txt;
paste -d '\n' ${g}_headers.txt ${g}_peptides.txt > ${g}_final.fasta
rm ${g}_peptides.txt
rm ${g}_headers.txt
done



#continuous

module load enthought_python/7.3.2
module load prinseq-lite/0.20.4
for file in /data/users/UserName/UserDirectory/job_trials/*.gz; do gunzip ${file};done
for file in 1_* 2_* 3_*; do mv "$file" "${file/S*_/A11.fastq}"; done
for file in 21_* 24_* 14_*; do mv "$file" "${file/S*_/OC.fastq}"; done
for file in 15_* 20_*; do mv "$file" "${file/S*_/Lock.fastq}"; done
for file in 96_* 97_* 98_*; do mv "$file" "${file/S*_/Bead.fastq}"; done
for file in *.fastq; do prinseq-lite.pl -fastq ${file} -out_format 3;done
find . -type f ! -name "*good*" -exec rm -rf {} \;
for file in *good*.fastq; do grep -o -P 'TTCTCACTCT.{0,36}' ${file}> ${file}_hit.txt; done
find . -type f ! -name "*.txt" -exec rm -rf {} \;
for file in *_hit.txt; do sed "s/TTCTCACTCT//g" ${file} > ${file}_final_thirtysix.txt; done
for file in *prinseq_good*.txt; do mv "$file" "${file/prinseq_good*/dna_sequence}"; done
for file in /data/users/UserName/UserDirectory/job_trials/*dna_sequence; do python
/data/users/UserName/UserDirectory/translate.py ${file}>${file}_translated; done
find . -type f ! -name "*translated" -exec rm -rf {} \; done
find /data/users/UserName/UserDirectory/job_trials -type f -exec sed -i 's/*/X/g' {} \;
find /data/users/UserName/UserDirectory/job_trials/ -type f -exec sed -i '/.\{12\}/!d' {} \;
for file in *Bead*translated ; do cat "$file" >> Bead_background.txt && rm "$file" || break ;
done
rm *Bead*good*
for file in *translated ; do grep -vxFf *background.txt ${file}> ${file}_bead_filter.txt; done
for i in *bead_filter.txt; do
cat ${i} | sort | uniq -c | sort -nr >> ${i}_bead_ranked.txt;
done
for i in *dna_sequence_translated_bead_filter.txt_bead_ranked.txt; do
g=$(basename $i dna_sequence_translated_bead_filter.txt_bead_ranked.txt);
sed -e "s/[0-9]*//g" ${i} | sed -e "s/ //g" > ${g}_peptides.txt
awk -v g="$g" '{ print ">"g"_peptide_"NR"_"$1$0}' ${i} | awk '{print $1}' > ${g}_headers.txt;
paste -d '\n' ${g}_headers.txt ${g}_peptides.txt > ${g}_complete.txt
```

```
rm ${g}_peptides.txt
rm ${g}_headers.txt
done
cat *A11*_bead_filter.txt > A11_all.txt
cat *OC*_bead_filter.txt > OC_all.txt
cat *Lock*_bead_filter.txt > Lock_all.txt
for i in *all.txt; do
cat ${i} | sort | uniq -c | sort -nr >> ${i}_sorted_ranked.txt;
done
for i in *_sorted_ranked.txt; do
awk '($1 > 4 ) ' ${i}>${i}_five.txt;
done
for i in *_sorted_ranked.txt; do
awk '($1 < 5 ) ' ${i}>${i}_low_counts.txt
done
for i in *_all.txt_sorted_ranked.txt_five.txt; do
g=$(basename $i _all.txt_sorted_ranked.txt_five.txt);
sed -e "s/[0-9]*//g" ${i} | sed -e "s/ //g" > ${g}_peptides.txt
awk -v g="$g" '{ print ">"g"_peptide_"NR"_"$1$0}' ${i} | awk '{print $1}' > ${g}_headers.txt;
paste -d '\n' ${g}_headers.txt ${g}_peptides.txt > ${g}_final.fasta
rm ${g}_peptides.txt
rm ${g}_headers.txt
done
```
